# Anabolic and catabolic responses to different modes of exercise in patients with chronic kidney disease

**DOI:** 10.1101/2025.09.15.676226

**Authors:** Douglas W Gould, Luke A Baker, Thomas J Wilkinson, Nicholas Eastly, Robert U Ashford, Matthew Denniff, Matthew Graham-Brown, Alice C Smith, João L Viana, Andy Philp, Emma L Watson

**Affiliations:** Intensive Care National Audit and Research Centre, London, UK; Division of Respiratory Sciences, School of Medical Sciences College of Life Sciences, University of Leicester, Leicester, UK; Leicester Biomedical Research Centre, Leicester Diabetes Centre, University of Leicester, Leicester, UK; Leicester Orthopaedics, University Hospitals of Leicester, UK; Department of Cancer Studies, University of Leicester, UK; Division of Cardiovascular Sciences, School of Medical Sciences, College of Life Sciences, University of Leicester, Leicester, UK; Leicester Kidney Lifestyle Team, Department of Population Health Sciences, University of Leicester, Leicester, UK; Research Center in Sports Sciences, Health Sciences and Human Development, CIDESD, University Institute of Maia, ISMAI, Portugal; Centre for Healthy Ageing, Centenary Institute, Sydney, New South Wales, Australia; School of Sport, Exercise and Rehabilitation Sciences, University of Technology Sydney, Sydney, New South Wales, Australia

## Abstract

**Background:** Muscle wasting is a common complication in individuals with chronic kidney disease (CKD) and contributes to reduced physical function and poor clinical outcomes. While exercise is recommended for CKD patients, the molecular responses to different exercise modalities remain poorly understood. This study aimed to investigate the anabolic, catabolic, and myogenic responses of skeletal muscle to aerobic exercise (AE) and combined exercise (CE; aerobic plus resistance) in people with CKD.

**Methods:** Muscle biopsies were collected from participants in a 12-week randomized controlled trial of supervised exercise training, the ExTRA CKD trial. Samples were obtained at baseline, 24 hours after an initial bout of exercise (untrained), and 24 hours after the final training session (trained). Western blotting and RT-qPCR were used to assess changes in key markers of protein synthesis, degradation, and regeneration. To complement these data, *in vitro* experiments using mechanically stretched primary skeletal muscle cells from CKD and healthy control donors were used to explore the time course of anabolic signalling.

**Results:** *In vivo*, Akt phosphorylation was blunted following unaccustomed CE but significantly upregulated following training, indicating partial restoration of anabolic signalling. No change was observed in response to AE. Myostatin expression was significantly downregulated following both AE and CE in the untrained state, while Pax7 and myogenic gene expression were upregulated only in response to CE after training. *In vitro*, mechanical stretch induced significant phosphorylation of Akt and p70S6K in both CKD and control cells, with no group difference in Akt response and a trend toward faster return to baseline of p70S6K phosphorylation in CKD cells.

**Conclusion:** These findings demonstrate that CE, but not AE, induces beneficial anabolic and myogenic responses in skeletal muscle in CKD, and highlight the value of combining *in vivo* and *in vitro* models to explore temporal dynamics and mechanistic insight into muscle adaptation.

**What was known:**

- People with CKD often have impaired skeletal muscle responses to exercise, termed anabolic resistance, which may contribute to muscle wasting and poor outcomes.
- Resistance exercise activates anabolic and myogenic pathways in healthy muscle, but the extent of these responses in CKD is unclear.
- Limited studies have compared molecular responses to different exercise modalities in CKD, particularly combining in vivo and in vitro approaches.

**This study adds:**

- Combined aerobic–resistance exercise, but not aerobic exercise alone, restored aspects of anabolic signalling (Akt phosphorylation) in CKD skeletal muscle after training.
- Myogenic gene expression was upregulated following combined exercise but not aerobic exercise, indicating a modality-specific regenerative response.
- *In vitro*, primary muscle cells from CKD and healthy donors showed similar early anabolic responses to mechanical stretch, though CKD cells may return to baseline more rapidly.

**Potential Impact:**

- Supports inclusion of resistance-based modalities in exercise prescriptions for people with CKD to enhance muscle anabolic and regenerative responses.
- Suggests that exercise prescriptions for CKD should prioritise combined modalities to maximise muscle anabolic and regenerative potential.
- Provides mechanistic evidence to inform tailored exercise interventions aimed at preserving muscle health in CKD patients.

## Introduction

Chronic kidney disease (CKD) is a public health emergency that has a global prevalence of approximately 9.1% [1]. Muscle wasting and a reduction in physical function are common at all stages of CKD, independent of age, [2] and are associated with poor clinical outcomes with one study showing that diagnosed sarcopenia increased the risk of all-cause mortality by 3-fold [3]. Furthermore, people at all stages of CKD have very high levels of physical inactivity [4], also an independent risk factor for all-cause mortality [5].

Although regular physical activity is strongly recommended for individuals with CKD, and formal exercise guidelines have recently been published for this population [6], structured exercise programmes remain underutilised. To maximise the benefits of exercise training, it is crucial to understand the underlying molecular mechanisms, particularly how different types of exercise affect skeletal muscle homeostasis. This understanding could guide the development of adjunctive strategies, such as targeted nutritional or pharmacological interventions. While a few human studies are beginning to provide data in this area [7–9], our understanding in the CKD population in particular remains limited.

Skeletal muscle mass is maintained by a dynamic balance between protein synthesis and degradation. Anabolic signalling through the Akt/mTOR pathway plays a central role in promoting muscle protein synthesis, while catabolic pathways, including the ubiquitin–proteasome system and myostatin signalling, regulate protein breakdown. In CKD, disturbances in this balance are common, often exacerbated by systemic factors such as insulin resistance, inflammation, and metabolic acidosis [10–12]. Impairments in anabolic signalling, particularly reduced phosphorylation of Akt and downstream targets, have been observed in animal models of CKD and are associated with muscle atrophy and impaired regeneration. Resistance exercise is known to activate these anabolic pathways in healthy individuals, but the extent to which this occurs in people with CKD remains under investigation.

Our previous work has shown that in CKD, the expected anabolic and mitochondrial responses to resistance exercise may be blunted, consistent with the concept of anabolic resistance [7, 9]. Specifically, we demonstrated attenuated activation of the insulin signalling pathway and limited upregulation of myogenic markers in CKD patients after a single bout of resistance exercise. However, following a period of training, some of these responses appeared to normalise. This finding has yet to be replicated.

Building on this previous work, the aim of this study was to investigate the anabolic and catabolic responses of skeletal muscle to both unaccustomed and accustomed exercise in individuals with CKD, using muscle biopsies collected from the previously published ExTra CKD randomised controlled trial [13]. In addition to analysing responses to aerobic and combined exercise training, we used an *in vitro* stretch model to better understand the temporal dynamics of exercise-induced molecular signalling in primary skeletal muscle cells from CKD and healthy donors. Together, these data provide a comprehensive view of how CKD muscle responds to exercise and highlight important considerations for optimising interventions aimed at preserving or improving muscle health in this population.

## Materials and methods

This study is a secondary analysis of the previously reported randomised controlled trial, the ‘ExTRA CKD’ trial [13] complemented by additional mechanistic *in vitro* investigations using primary cells obtained from participants in the ‘Explore CKD’ trial.

### The ExTRA CKD trial

In brief, 54 non-dialysis patients with CKD stages 3b–5 were enrolled in a 12-week, thrice-weekly supervised exercise intervention, randomised to either aerobic exercise (AE) or combined exercise (CE). Each participant served as their own control through a six-week run-in period prior to randomisation. The AE intervention consisted of circuit-based aerobic activities such as treadmill walking or running, cycling, and rowing. The CE group undertook the same aerobic training, with the addition of resistance exercises performed during two of the three weekly sessions. Participants aimed to complete 30 minutes of moderate-intensity exercise at 70–80% of their heart rate maximum, determined by a maximal exercise tolerance test. For the resistance component, participants performed 3 sets of 12–15 repetitions of leg extensions at 70% of their estimated one-repetition maximum.

Recruitment occurred between 16^th^ December 2013 and 30^th^ April 2016, with all interventions completed by October 2016. Ethical approval was obtained from the UK National Research Ethics Committee (Ref: 13/EM/0344), and all participants provided written informed consent. The trial was conducted in accordance with the Declaration of Helsinki and registered with ISRCTN (No. 36489137). The study was registered with the ISRCTN (no. 36489137).

A subset of participants (AE, n = 10; CE, n = 9) consented to skeletal muscle biopsies of the vastus lateralis, as part of the ExTRA CKD trial, using a needle biopsy technique previously described [9]. Samples were collected in a fasted state at baseline, 24 hours after the first training session (“untrained”), and 24 hours after the final session (“trained”). Following removal of visible fat, biopsies were snap-frozen in liquid nitrogen and stored for later analysis.

### The Explore CKD Trial

A single skeletal muscle biopsy was obtained from six individuals with CKD and eight age- and sex-matched non-CKD controls as part of the ‘Explore CKD’ trial. CKD participants were recruited from outpatient clinics at Leicester General Hospital, UK, between 1^st^ January 2016 and 2^nd^ November 2020, and biopsies were collected using the needle biopsy technique. Healthy controls (HC), with no significant medical history, were recruited from orthopaedic theatre lists during procedures for benign tumour removal; their muscle biopsies were obtained via the open biopsy technique. The study received ethical approval from the UK National Research Ethics Committee (Ref: 15/EM/0467), and all participants provided written informed consent. The trial was conducted in accordance with the Declaration of Helsinki and registered with ISRCTN (No. 18221837).

### Molecular biology techniques

#### Western Blotting

15-20mg/ww muscle tissue was homogenised in 18μl/mg RIPA buffer (Sigma, UK) supplemented with 1% v/v phosphatase inhibitor-3 (Sigma Aldrich, UK), rotated for 90 minutes at 4°C, and centrifuged at 13,000rpm for 15 minutes at 4°C. The supernatant was collected and protein concentration determined by the Bio-Rad Protein Assay (BioRad, UK,). The pellet was retained for 14kDa actin fragment analysis [14]. Lysates were subjected to SDS-PAGE using 10-12% gels on a mini-Protean Tetra system (Bio-Rad, UK). Proteins were transferred onto nitrocellulose membranes, blocked for 1h with Tris-buffered saline with 5% (w/v) skimmed milk and 0.1% (v/v) Tween-20 detergent. Membranes were incubated with the primary antibody overnight. Antibodies to determine p-AktSer^473^ (1:2000), p-P70S6KThr^389^ (1:500) were obtained from Cell Signalling Technologies (Danvers, MA, USA) and AC40 Actin clone (1:500; Sigma Aldrich, UK) for analysis of the 14kDa fragment. The AC40 clone antibody also recognises the 42kDa fragment, which was used as a loading control with a much shorter exposure to prevent over-exposure. For all other proteins, GAPDH was used as a loading control. Horseradish Peroxidase (HRP) linked secondary anti-mouse/rabbit secondary antibodies (Dako, Aglient, UK) were used at 1:1500 for 2h at room temperature. Blots were visualised using ECL Reagents (Geneflow, UK) captured using a ChemiDoc MP imager (Bio-Rad).

#### Quantitative RT-PCR

Total RNA was isolated from 15-20mg/ww muscle tissue using TRIzol® (Invitrogen, UK) and reversed transcribed to cDNA using an AMV reverse transcription system (Promega, WI, USA). Primers and probes and internal controls were supplied as Taqman gene expression assays (Applied Biosystems, UK). MAFbx Hs00369714_m1, MuRF-1 Hs00822397_m1, Myostatin Hs00976237_m1, activin receptor IIB, Hs00155658_m1, Myogenin Hs01072232_m1, MyoD Hs02330075_g1, Myf5 Hs00929416, Pax 7 Hs00242962, and 18S Hs99999901 which was used as a housekeeping gene with a coefficient of variation of 2.18%. All reactions were carried out in a 20μl volume, 1μl cDNA, 10μl 2X Taqman Fast Mastermix, 8μl water, 1μl primer/probe on an Agilent Biosystem’s Light Cycler with the following conditions, 95°C 15 seconds, followed by 40X at 95°C for 15 seconds and 60°C for 1 min. The Ct values from the target gene were normalised to the Ct values for the house-keeping gene (18s, expression of which was found to be stable in these conditions, data not shown) and the expression levels calculated according to 2^−ΔΔCt^ method, which by definition sets the baseline value to 1.0.

#### Satellite cell isolation procedure

Satellite cells were isolated and propagated as previously described [15]. Muscle tissue was washed three times in HamsF10 (containing 1% penicillin streptomycin and 1% Gentamycin), minced into small fragments and enzymatically digested in two incubations with collagenase IV (1mg/mL), BSA (5mg/mL) and trypsin (500μl/mL) at 37°C with gentle agitation. The resultant supernatant was strained through a 70μm nylon filter and centrifuged at 800 g for 7 minutes. The cells were washed in Hams F10 with 1% penicillin streptomycin and pre-plated on uncoated 9cm2 petris in 3mL growth medium (GM; Hams F10 Glutamax, 20% FBS, 1% penicillin streptomycin, 1% fungazone) for 3h. The cell suspension was then moved to collagen I coated 25 cm2 flasks and kept in standard culture environmental conditions (37°C, 5 % CO2). For the expansion of satellite cell populations, cells were grown to approximately 70% confluence in GM that was changed every other day. Cells were subsequently trypsinised (Sigma-Aldrich, UK) and counted using the trypan blue exclusion method.

#### Cell Culture and treatments

Myoblasts were seeded on 6-well Flexcell culture plates, and cyclic multiaxial stretch applied using a Flexcell Fx3000 system set to 2 second sine wave stretch with 4 second release resulting in an 18% maximum stretch for 30 minutes at 37°C. Cells were harvested in 200μl lysis buffer/well RIPA buffer (Sigma, UK) supplemented with 1% v/v phosphatase inhibitor-3 (Sigma Aldrich, UK) prior to stretch, immediately post and at 1,3,7 and 24h post stretch. Protein concentration was determined by the Bio-Rad Protein Assay, and resulting lysates were stored for subsequent western blotting analysis.

#### Statistics

The distribution of the data was assessed using the Shapiro-Wilks test. Variables that were not normally distributed were log-transformed for analysis, and back transformed for presentation. A primary analysis was performed for each variable using repeated measures ANOVA for biopsy time-point (baseline, ‘untrained’, and ‘trained’) and group allocation (AE and CE). In addition, paired t-tests were performed between specific time points (baseline - untrained, baseline - trained, and untrained - trained) following AE and CE.

## Results

### Participant characteristics

Characteristics of the patients included in this study from the ExTRA CKD and Explore CKD trials can be found in Tables 1 and 2, respectively. The groups were well matched for all characteristics, with healthy controls demonstrating eGFR values within the normal range.

**Table 1.**
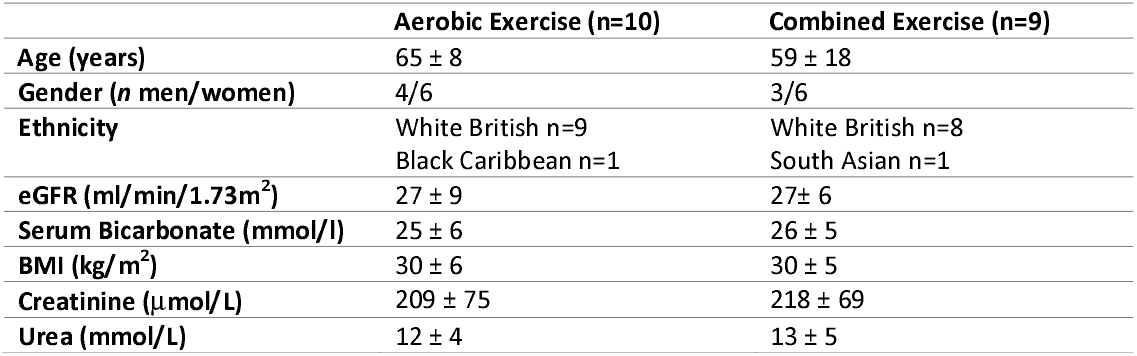
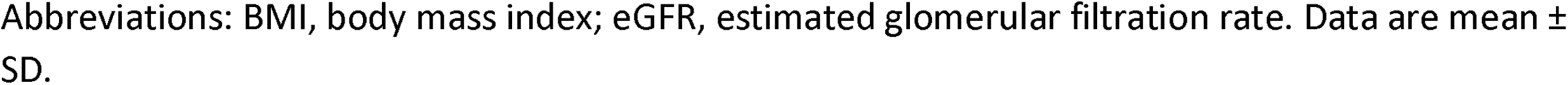
Participant characteristics.

|  | Aerobic Exercise (n=10) | Combined Exercise (n=9) |
| --- | --- | --- |
| Age (years) | 65 ± 8 | 59 ± 18 |
| Gender (n men/women) | 4/6 | 3/6 |
| Ethnicity | White British n=9<br>Black Caribbean n=1 | White British n=8<br>South Asian n=1 |
| eGFR (ml/min/1.73m<sup>2</sup>) | 27 ± 9 | 27 ± 6 |
| Serum Bicarbonate (mmol/l) | 25 ± 6 | 26 ± 5 |
| BMI (kg/m<sup>2</sup>) | 30 ± 6 | 30 ± 5 |
| Creatinine (μmol/L) | 209 ± 75 | 218 ± 69 |
| Urea (mmol/L) | 12 ± 4 | 13 ± 5 |

Abbreviations: BMI, body mass index; eGFR, estimated glomerular filtration rate. Data are mean ± SD.

**Table 2.** Skeletal muscle biopsy donors for primary cell culture establishment.

|  | Healthy Controls (n=8) | CKD Patients (n=6) |
| --- | --- | --- |
| **Age (years)** | 61 ± 14 | 63 ± 6 |
| **Gender (*n* men/women)** | 4/4 | 2/3 |
| **Ethnicity** | White British n=8 | White British n=6 |
| **eGFR (ml/min/1.73m<sup>2</sup>)** | 25.0 ± 4.7 | 82.0 ± 9.9 |

Abbreviations: eGFR, estimated glomerular filtration rate. Data are mean ± SD.

### The anabolic response to exercise in CKD

Changes in phosphorylation of Akt and P70S6K are shown in Figure 1. Akt phosphorylation is well documented to increase in the hours following exercise, and is a key contributor to the anabolic response to exercise [16]. No significant changes in Akt phosphorylation from baseline were observed at any time point following AE. There was also no change in Akt phosphorylation from baseline following an acute bout of CE in the untrained state (P=0.48, Figure 1A-D). However, following 12-weeks of CE training, Akt phosphorylation was upregulated 118% from baseline (P=0.02) and was significantly higher than the response before training (P=0.02). There was no change in the phosphorylation status of P70S6K in response to an acute bout of AE or CE either before or after training (Figure 1E-H).

**Figure 1.**
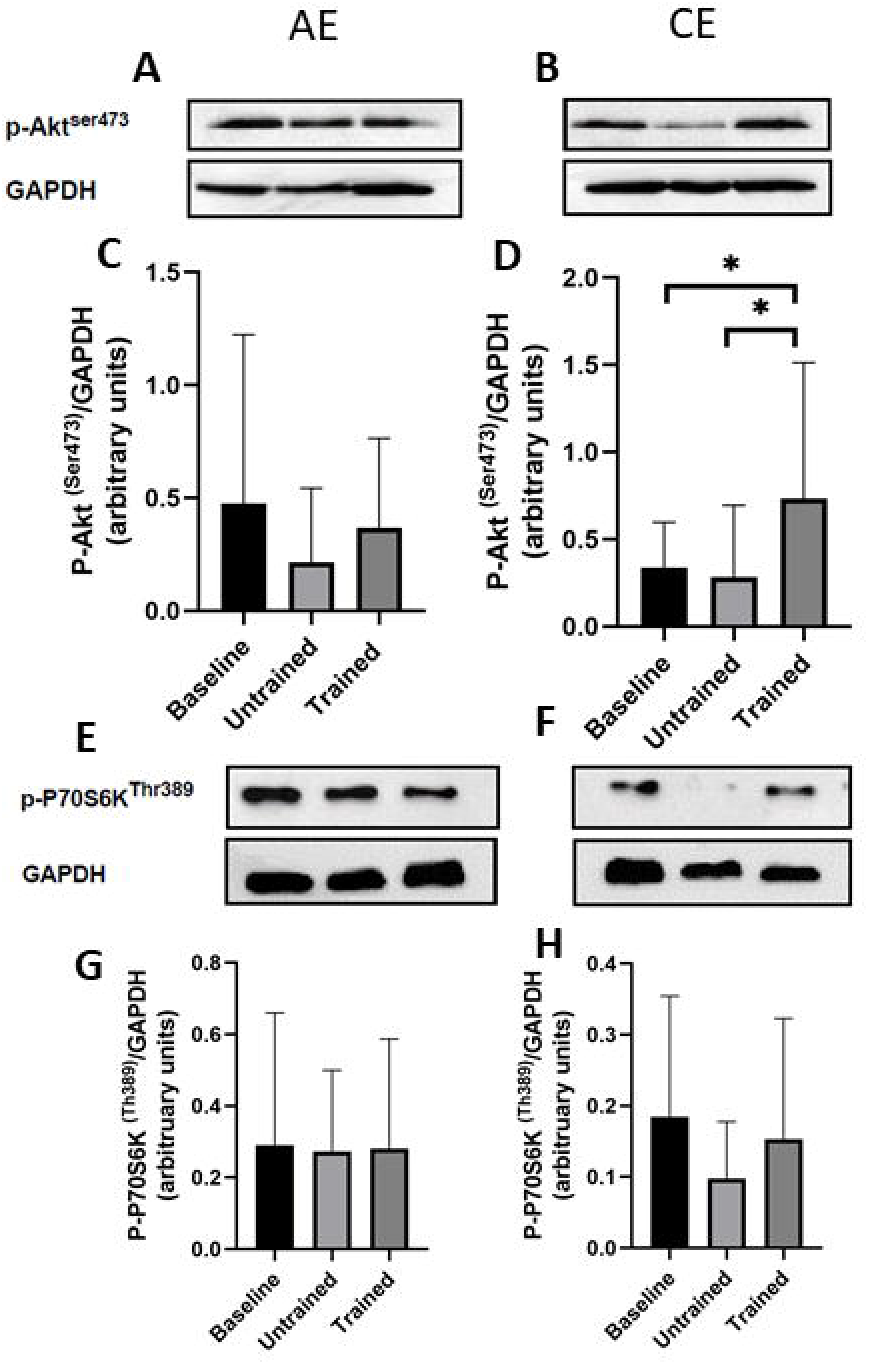
Changes in phosphorylation of Akt and P70S6K in skeletal muscle biopsies in response to unaccustomed and accustomed bouts of AE or CE. People with CKD undertook 12 weeks of either AE or CE training. Vastus lateralis muscle biopsies were collected at baseline, 24h after a bout of unaccustomed exercise (untrained) and 24h after a bout of accustomed exercise (Trained) in both groups. A) shows a representative western blot image for P-AktSer^473^ at baseline, untrained and trained time points with GAPDH that was used as a loading control in those people randomised to AE, and B) in those randomised CE. C-D) Histograms displaying densitometric data. E) shows a representative western blot image for P-70S6KThr^389^ at baseline, untrained and trained time points with GAPDH that was used as a loading control in those people randomised to AE and F) in those randomised to CE. For both proteins, AE n= 9, CE n=8. Data are presented as mean±SD. * denotes P<0.05. Abbreviations: AE, aerobic exercise, CE, combined exercise.

### The catabolic response to exercise in CKD

The presence of the 14kDa fragment within biopsy samples can be found in Figure 2. The 14kDa is a cleavage product of actin and myosin and has been used as a biomarker of skeletal muscle catabolism [14]. The abundance of the 14 kDa fragment remained unchanged at all time points following AE. In contrast, an acute bout of unaccustomed CE prior to training resulted in a 253% increase in the 14 kDa fragment relative to baseline (P = 0.04); this response was no longer evident after 12 weeks of CE training, with levels returning to baseline (P = 0.68 vs baseline). MuRF-1 and MAFbx are two muscle specific E3 ligases, commonly used as markers of ubiquitin-proteasome system activity [17]. Repeated measures ANOVA revealed there was no effect of acute AE or CE on MuRF-1 mRNA expression (P=0.46; Figure 3A-B) or MAFbx mRNA expression (P=0.70; Figure 3A-B). In contrast to the pattern of expression seen with the E3 ligases, myostatin, a potent catalyst for skeletal muscle protein wasting [18], was significantly down regulated from baseline following an acute bout of AE in both the untrained (1.5-fold reduction; P=0.01; Figure 3C) and trained (1.2-fold reduction; P=0.04) conditions. In response to an acute bout of CE in the untrained state, myostatin mRNA expression decreased by 5-fold (P = 0.003; Figure 3D). However, myostatin mRNA expression in response to acute CE after training was significantly elevated above the untrained response (P=0.007; Figure 3D). No changes were seen in any condition in the expression of the myostatin receptor ActIIRB (P>0.05; Figure 3C-D).

**Figure 2.**
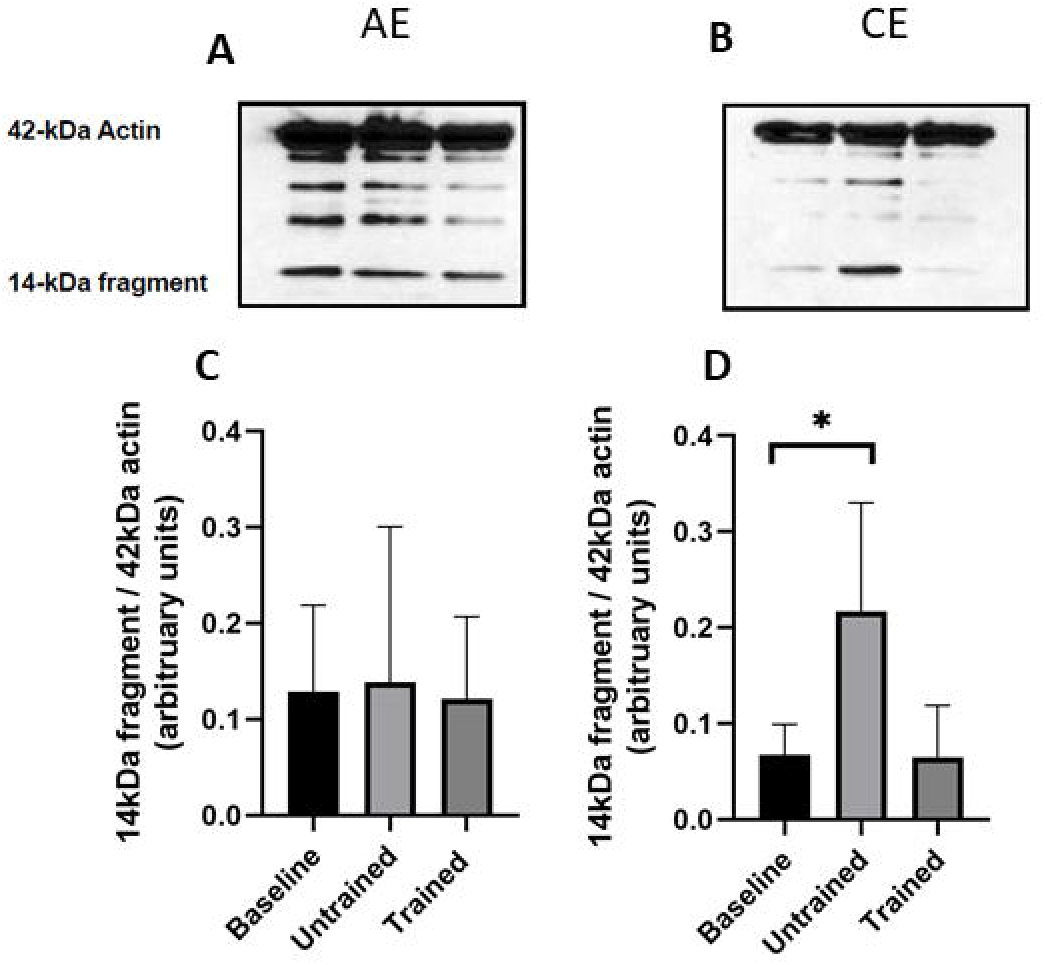
Abundance of the 14kDa actin fragment in skeletal muscle biopsies in response to unaccustomed and accustomed bouts of AE or CE. People with CKD undertook 12 weeks of either AE or CE training. Vastus lateralis muscle biopsies were collected at baseline, 24h after a bout of unaccustomed exercise (untrained) and 24h after a bout of accustomed exercise (Trained) in both groups. A) shows a representative full western blot image that are labelled to show the 42kDa and the 14kDa actin fragments from people within the AE group and B) CE group. Histograms in C-D show densitometric data. AE n= 6, CE n=5. Data are presented as mean±SD. * denotes P<0.05. Abbreviations: AE, aerobic exercise, CE, combined exercise.

**Figure 3.**
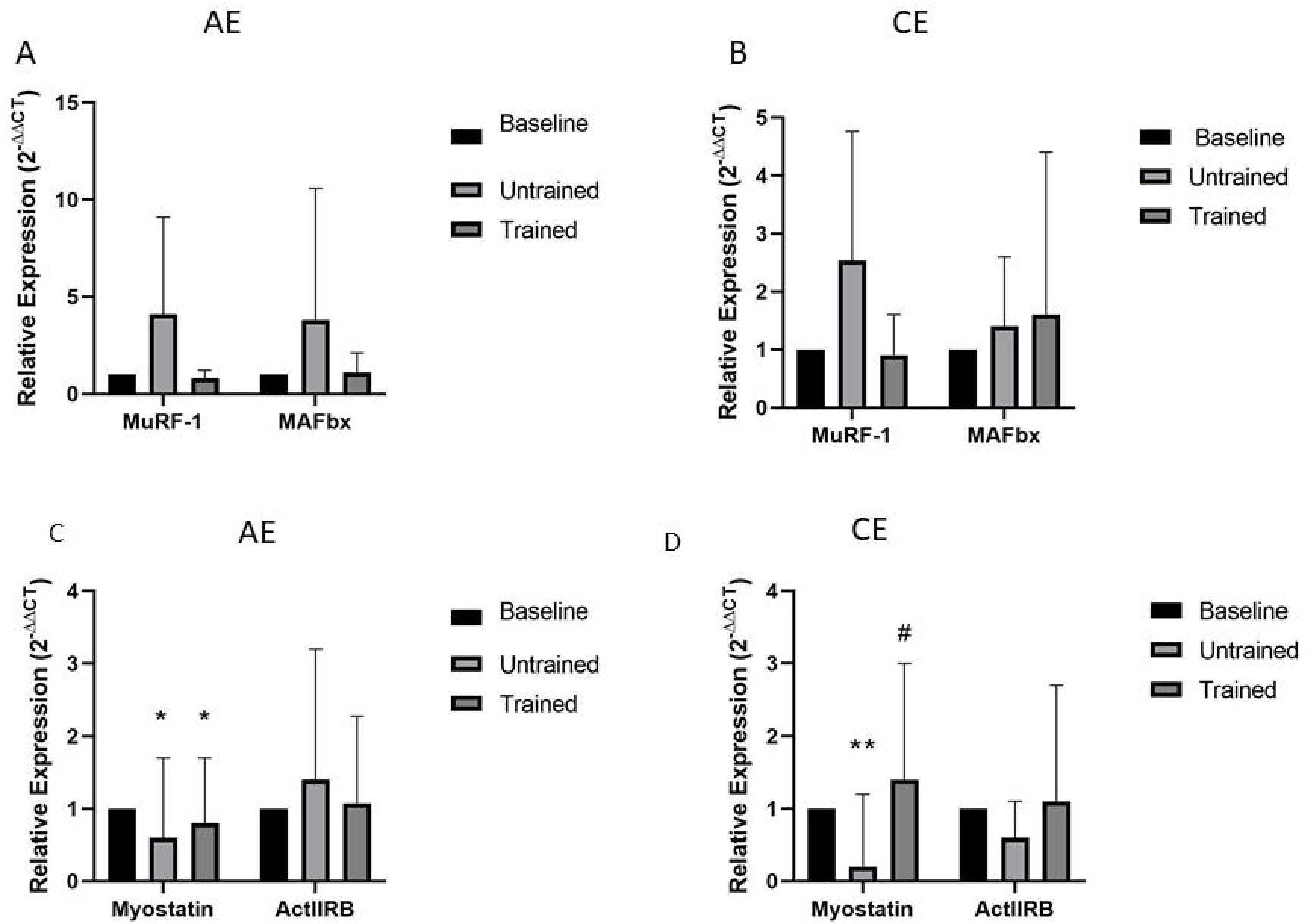
Changes in mRNA expression of genes related to atrophy processes in response to unaccustomed and accustomed bouts of AE or CE. People with CKD undertook 12 weeks of either AE or CE training. Vastus lateralis muscle biopsies were collected at baseline, 24h after a bout of unaccustomed exercise (untrained) and 24h after a bout of accustomed exercise (Trained) in both groups. mRNA expression of MuRF-1 and MAFbx were analysed by RT-PCR in the A) AE group, and B) CE group. Myostatin and ActIIRB were analysed by RT-PCR in the C) AE group, and D) CE group. Expression is displayed as relative change from baseline according to 2−ΔΔCt method and normalized to 18S. Data are presented as mean±SD. MuRF-1 AE n= 9, CE n=8; MAFbx AE n=8 CE = 10; Myostatin AE=10, CE=9; AC2BR AE=10, CE=9. * denotes P<0.05 vs baseline, ** denotes P<0.01 vs baseline, # denotes P<0.05 vs untrained. Abbreviations: AE, aerobic exercise; CE, combined exercise.

### The myogenic response to exercise in CKD

The myogenic factors Pax7, MyoD, myogenin and Myf5 coordinate the process of skeletal muscle repair and regeneration [19]. Changes in the mRNA of these genes can be found in Figure 4. There was no change in the mRNA expression of any of these factors at any time point following AE (P<0.05; Figure 4A). There was also no change in the mRNA expression of Myogenin or Myf5 following CE exercise at any timepoint (P<0.05; Figure 4B). We did observe a near significant 2-fold decrease in MyoD mRNA expression following an acute bout of CE exercise in the untrained state (P=0.05; Figure 4B), a response that significantly increased following training (P=0.01 trained vs untrained condition; Figure 4B). Finally, Pax7 mRNA expression in response to a bout of CE was significantly greater following training compared to the pre-training response (P = 0.025; Figure 4B).

**Figure 4.**
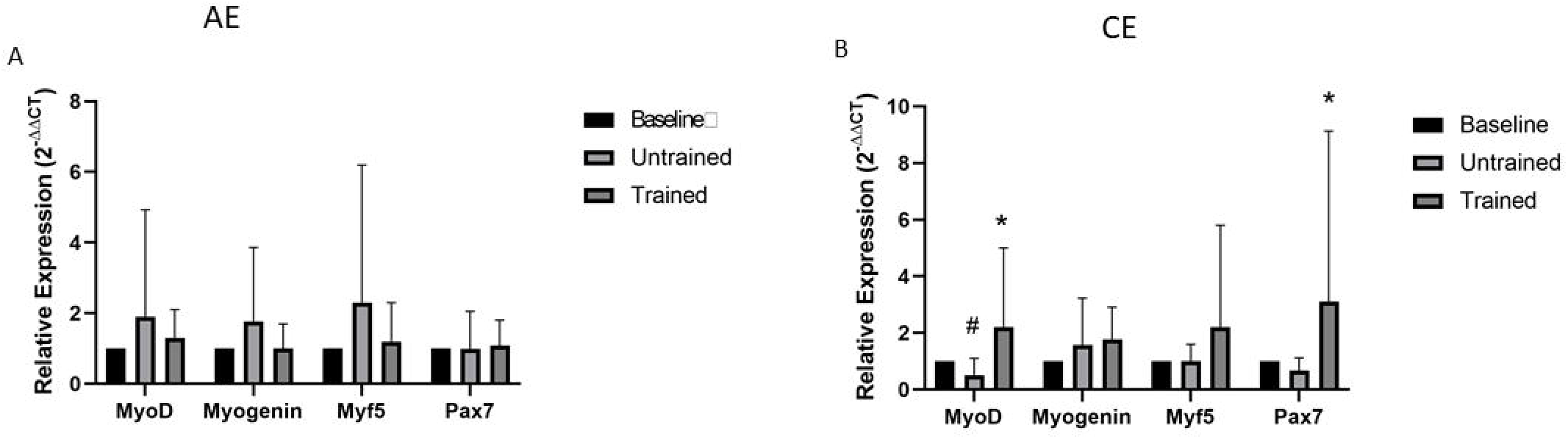
Changes in mRNA expression of genes related to myogenesis in response to unaccustomed and accustomed bouts of AE or CE. People with CKD undertook 12 weeks of either AE or CE training. Vastus lateralis muscle biopsies were collected at baseline, 24h after a bout of unaccustomed exercise (untrained) and 24h after a bout of accustomed exercise (Trained) in both groups. mRNA expression of genes within the myogenic pathways were analysed by RT-PCR in the A) AE group and B) CE group. Expression is displayed as relative change from baseline according to 2−ΔΔCt method and normalized to 18S. Data are presented as mean±SD. MyoD AE n=10, CE n=9; Myogenin AE n=10, CE n=9, Myf5 AE n=8, CE n=9; Pax 7 AE n=10, CE n=8.* denotes P<0.05 vs untrained. # denotes P=0.05 vs baseline. Abbreviations: AE, aerobic exercise; CE, combined exercise.

### *In-Vitro* response to cell stretch

In order to examine more closely the changes in phosphorylation of key proteins controlling protein synthesis events, we mechanically stretched cells from CKD and HC donors *in vitro*, and collected the cells at regular intervals post stretch up to 24h. This provides key information about the immediate changes that we were not able to capture in the *ex vivo* biopsy study. Mechanical stretch induced a significant increase in Akt phosphorylation relative to baseline in both CKD and HC groups immediately post-stretch (CKD: +5651%, P = 0.012; HC: +3437%, P = 0.028; Figure 5A,C,D), and this elevation persisted at 1-hour post-stretch (CKD: +61%, P = 0.012; HC: +1472%, P = 0.028). At 3-hours post-stretch, Akt phosphorylation remained significantly elevated in the HC group (+388%, P = 0.046), but this was not seen in the CKD group (+48%, P = 0.31). By 7- and 24-hours post-stretch, phosphorylation levels had returned to baseline in both groups (Figure 5A). Repeated measures ANOVA revealed no significant interaction effect, indicating that the response to the stretch protocol did not differ between the CKD and HC groups (P = 0.84).

**Figure 5.**
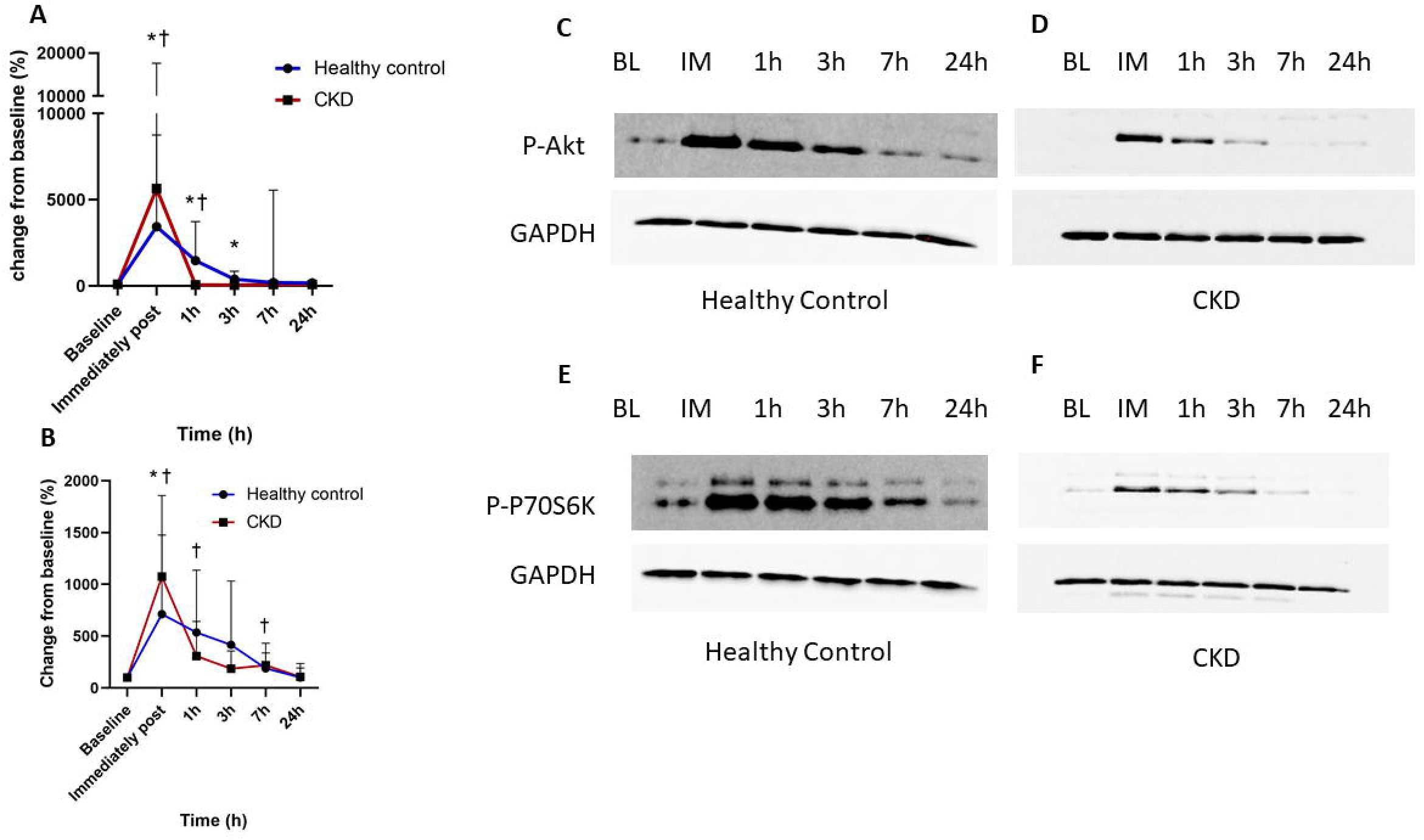
The effect of 18% cyclic stretch on the phosphorylation of Akt and P70S6K in primary skeletal muscle myotubes from healthy control or CKD donors. Myotubes were stretched on 6-well Flexcell culture plates for 30 minutes using a Flexcell Fx3000 system set to 2 second sine wave stretch with 4 second release. Cells were harvested for western blotting prior to stretch (baseline), immediately post, and at 1,3,7 and 24h post stretch for the analysis of P-Akt and P-P70S6K. A) Time course of % change from baseline of P-Akt, n=8 healthy control, n=6 CKD, and B) P-P70S6K n=7 healthy control, n=5 CKD. Representative western blot images of C) P-Akt normalised to GAPDH in a healthy control donor and D) P-Akt normalised to GAPDH in a CKD donor E) P-P70S6K normalised to GAPDH in a healthy control donor and F) P-P70S6K normalised to GAPDH in a CKD donor. Data are presented as mean±SD. * Denotes P<0.05 vs baseline in HC. † denotes P<0.05 vs baseline in CKD. Abbreviations: BL, Baseline; CKD, chronic kidney disease; h, hour; IM, immediately post stretch.

A similar response was seen for P70S6K phosphorylation (Figure 5B,E,F). P70S6K phosphorylation was seen to significantly increase from baseline immediately post stretch in both groups (CKD: +1076%, P=0.014; HC +712%, P=0.038). In CKD cells, p70S6K phosphorylation remained significantly elevated at 1-hour post-stretch (+306%, P = 0.047), with a strong trend toward an increase at 3 hours (+185%, P = 0.052), and a significant upregulation observed again at 7-hours post-stretch (+219%, P = 0.004) In contrast, no further time points showed significant increases from baseline in the HC group, likely due to high inter-individual variability. A repeated measures ANOVA showed a near interaction effect, but this just fell short of significance (P=0.052), indicating broadly similar responses to the stretch regime between the two groups.

## Discussion

This study offers new insights into the molecular responses to aerobic and combined exercise in individuals with CKD, while also supporting findings from our previous research in a different cohort and exercise modality [9]. Evidence suggests that individuals with CKD do not always exhibit the expected physiological adaptations to exercise. For instance, several studies have reported that aerobic training, typically associated with improved exercise capacity in the general population, does not consistently lead to increases in VO_2peak_ in CKD patients [20–22] a finding linked to a blunted activation of mitochondrial biogenesis pathways [7]. In addition, following a bout of resistance exercise training it was reported that the expected increases in activation of proteins within the insulin signalling pathway were largely absent, but that this response was restored after training [9] a finding that we have replicated here. Understanding the molecular responses that underpin these physiological adaptations to exercise will enable us to design adjunct therapies to maximise the benefits that patients receive from exercise training.

Skeletal muscle homeostasis is maintained through a finely regulated balance between protein synthesis and protein degradation. In CKD, insulin resistance has been implicated in the muscle wasting commonly observed in patients [10], which can be seen in reduced phosphorylation of the key proteins in the Akt/mTOR pathway [11, 12], an abnormality that can be reversed by exercise training in CKD mice [11]. It is well established that resistance exercise strongly stimulates pathways involved in protein synthesis, with activity remaining elevated for up to 48 hours post-exercise [23]. Our previous work [9] indicated that individuals with CKD exhibit a blunted anabolic response to an acute bout of exercise in the untrained state, consistent with the presence of anabolic resistance. Importantly, this response was restored following a period of training. The current study confirms these findings, in participants undergoing combined exercise training, there was no detectable change in Akt phosphorylation in response to exercise before training. However, after 8 weeks of training, Akt phosphorylation was significantly elevated above baseline, which could indicate a partial restoration of anabolic signalling. No change in Akt or P70S6K phosphorylation was observed in the aerobic training group, which aligns with previous evidence that aerobic exercise does not robustly activate this pathway [24].

A major limitation of muscle biopsy studies is the limited temporal resolution they offer, often providing only a snapshot of complex, dynamic processes. In this study, muscle biopsies were collected 24 hours post-exercise, meaning that early molecular responses may have been missed due to the chosen sampling timepoint. To address this gap, we conducted *in vitro* stretch experiments using primary human skeletal muscle cells from both healthy controls and individuals with CKD, cells known to retain their *in vivo* phenotype [15] and matched for age, gender and physical activity levels (all had low physical activity levels). These experiments demonstrated that mechanical stretch significantly increased phosphorylation of both Akt and p70S6K in both groups, between immediately post stretch and 7h post stretch, with phosphorylation levels returning to baseline at 24h, replicating the *in vivo* data at this time point. Interestingly, no significant difference was observed between CKD and control cells in their Akt phosphorylation response to stretch. Although a strong trend toward a group difference in p70S6K phosphorylation was observed, it narrowly missed statistical significance; CKD cells exhibited a higher initial response to stretch, which appeared to return to baseline more rapidly than in healthy controls, where the response was more sustained. This may be due to insufficient statistical power, especially given the high variability observed in the data. Notably, each replicate represented a different donor. Primary cultures from different individuals may respond variably to stretch, therefore future studies using multiple replicates from the same donor may help clarify the consistency and magnitude of individual responses. Integration of the *in vivo* and *in vitro* data suggests that individuals with CKD are capable of mounting an anabolic response to exercise; however, this response appears to be relatively transient. Conducting an *in vitro* ‘training’ model using repeated bouts of mechanical stretch could offer valuable insights into how chronic loading influences the durability and magnitude of the anabolic signalling response in CKD patients.

An essential response to muscle damage is the activation of satellite cells, the resident muscle stem cell population which, upon activation, migrate to the site of injury to support muscle repair and regeneration. In the AE group, there were no significant changes in MyoD, Myf5, Myogenin, or Pax7 mRNA expression across time points, suggesting that aerobic exercise alone does not robustly stimulate myogenic gene expression in this population. In contrast, the CE group showed a more pronounced response, MyoD and Myogenin expression were significantly upregulated 24 hours after an accustomed bout of CE compared to the untrained condition. Pax7 expression was significantly higher post-training compared to the untrained state, indicating enhanced satellite cell proliferation following repeated combined exercise. Given that resistance-based exercise, which forms part of the CE intervention, is more likely to induce muscle damage than aerobic exercise alone, it is plausible that regenerative processes, including satellite cell-mediated repair, are more strongly upregulated in response to CE than AE.

Several limitations should be acknowledged. The timing of muscle biopsy collection restricted to a single time point 24 hours post-exercise limits the conclusions that can be drawn to a narrow temporal window. As such, early molecular events that may have occurred in the immediate hours following exercise could have been missed. This limitation was a key rationale for conducting the complementary *in vitro* study, which allowed for a more detailed examination of the temporal profile of anabolic signalling. It must be highlighted that whilst a good *in vitro* model, mechanical stretch of human primary muscle cells does not fully recapitulate exercise *in vivo*. It does not capture the complex interactions between hormonal changes, neural input or immune responses that can all affect the molecular response within the muscle. 2D cultured primary cells can also lose aspects of their tissue-level organisation and environmental context over time. Factors such as extracellular matrix composition, cell–cell interactions, and 3D architecture, which influence signal transduction *in vivo*. To better capture the dynamics of molecular responses to exercise in future *in vivo* studies, multiple biopsy time points across the acute post-exercise period should be considered.

In conclusion this study provides novel insight into the molecular responses of skeletal muscle to different modes of exercise in individuals with CKD. Using both in vivo muscle biopsy data and complementary in vitro models, we demonstrate that combined exercise training, but not aerobic training alone, can restore aspects of anabolic signalling, namely Akt phosphorylation, in skeletal muscle, supporting the notion that resistance-based modalities are necessary to elicit beneficial molecular adaptations in this population and confirming our earlier work. While the in vitro stretch model helped clarify the temporal dynamics of early signalling events, it also highlighted the complexity of translating findings across experimental models. Our findings suggest that individuals with CKD are capable of mounting an anabolic and myogenic response to exercise, but that this response may be blunted or short-lived without repeated loading. The observed changes in catabolic and regenerative markers further support the importance of exercise modality in shaping muscle adaptation and further validating the role of exercise training in reversing anabolic resistance in CKD. Future work should focus on longitudinal models, both *in vitro* and *in vivo*, to better understand how repeated loading influences anabolic signalling, and how these molecular responses relate to clinically meaningful improvements in muscle mass and function in CKD.

## Acknowledgements

Authors declare no conflict of interest. This is a summary of independent research funded by the Leicester Hospitals Charity Kidney Care Appeal and the Stoneygate Trust, supported by the National Institute for Health and Care Research (NIHR) Leicester Biomedical Research Centre (BRC). The views expressed are those of the author(s) and not necessarily those of the NIHR or the Department of Health and Social Care. For the purpose of open access, the author has applied a Creative Commons Attribution license (CC BY) to any Author Accepted Manuscript version arising from this submission.

